# Influenza virus antibodies inhibit antigen-specific *de novo* B cell responses in mice

**DOI:** 10.1101/2024.04.12.589218

**Authors:** Eileen Goodwin, James S. Gibbs, Jonathan W. Yewdell, Laurence C. Eisenlohr, Scott E. Hensley

## Abstract

Antibody responses to influenza vaccines tend to be focused on epitopes encountered during prior influenza exposures, with little production of *de novo* responses to novel epitopes. To examine the contribution of circulating antibody to this phenomenon, we passively transferred a hemagglutinin (HA)-specific monoclonal antibody (mAb) into mice before immunizing with whole inactivated virions. The HA mAb inhibited *de novo* HA-specific antibodies, plasmablasts, germinal center B cells, and memory B cells, while responses to a second antigen in the vaccine, neuraminidase (NA), were uninhibited. The HA mAb potently inhibited *de novo* antibody responses against epitopes near the HA mAb binding site. The HA mAb also promoted IgG1 class switching, an effect that, unlike the inhibition of HA responses, relied on signaling through Fc-gamma receptors. These studies suggest that circulating antibodies inhibit *de novo* B cell responses in an antigen-specific manner, which likely contributes to differences in antibody specificities elicited during primary and secondary influenza virus exposures.

## Introduction

Despite decades of research, influenza vaccines remain less effective than many other human vaccines (Osterholm et al., 2012). Influenza vaccines are updated regularly with the goal of eliciting antibodies against novel epitopes on new circulating strains. However, these vaccines often fail to elicit strong *de novo* antibody responses and instead boost responses to previously encountered influenza strains. This phenomenon, termed “original antigenic sin” (OAS) (Francis, 1960), presents a substantial barrier to improving influenza vaccine effectiveness. A better understanding of how pre-existing immunity affects the generation of *de novo* immune responses will allow for the rational design of better influenza vaccines.

During secondary immune responses, memory B cells rapidly differentiate into antibody-secreting cells (ASCs) (Mesin et al., 2020; Pape et al., 2011), leading to the production of high levels of class-switched antibody. Memory B cells elicited by initial antigen exposures produce the majority of long-lived serum antibody, while naïve B cells stimulated during second exposures appear to be inefficient at differentiating into ASCs (Schiepers et al., 2023) despite being abundant in secondary germinal centers (Mesin et al., 2020). It is unknown what components of immune memory inhibit naïve B cells from efficiently maturing into ASCs during secondary immune responses. One possibility is that circulating antibodies elicited by prior exposures prevent naïve B cells from becoming ASCs.

The capacity of circulating antibody to inhibit immune responses is well established. For example, passive administration of anti-D antibody is used to block *de novo* generation of anti-D antibodies in RhD-negative mothers, thereby preventing hemolytic disease of the newborn (Kumpel & Elson, 2001; Urbaniak & Greiss, 2000). Similarly, maternally-derived antibodies in neonates prevent the generation of protective immune responses to vaccines early in life (Siegrist, 2003; Voysey et al., 2017). These inhibitory effects are not unique to passively acquired antibody: the amount of pre-existing vaccine-reactive antibody is inversely correlated with the amount of antibodies elicited by influenza vaccines in humans (Zarnitsyna et al., 2015). The immunological basis of this apparent antibody inhibition of influenza vaccine responses is not well understood.

To determine how influenza virus antibodies affect *de novo* B cell responses, we, passively administered an anti-hemagglutinin (HA) monoclonal antibody (mAb) into mice and profiled its impact on the antibody response to whole inactivated influenza virus. We identified two main effects of the HA mAb: inhibition of *de novo* antibodies to epitopes near the HA mAb binding site and a broad enhancement of IgG1 against other HA epitopes farther away from the mAb binding site and non-HA proteins. These results show that circulating influenza virus antibodies can broadly impact *de novo* B cell responses and may influence the specificity of antibody responses during secondary influenza virus encounters.

## Results

### HA-specific antibody potently inhibits antibody responses to HA but not other viral antigens

We administered a mAb that binds to the Sb antigenic site (Caton et al., 1982) of H1N1 HA (H28-D14; IgG2a isotype) or a non-influenza virus IgG2a mAb to BALB/c mice prior to vaccination with whole inactivated virions of A/Puerto Rico/8/1934 H1N1 influenza virus. At the time of vaccination, the HA mAb was present in serum at approximately 0.5 μg/ml (**Supplementary Fig. 1A-B**). We measured vaccine-specific serum antibodies in these mice at 7, 14, and 28 days following vaccination. In the presence of the HA mAb, the overall antibody response to HA (as measured by ELISA with an anti-kappa light chain detection antibody) was greatly reduced but not eliminated (**Fig. 1A**). The corresponding antibody response to neuraminidase (NA), another protein antigen present in the vaccine, was mostly unaffected (**Fig. 1A**).

**Fig. 1:**
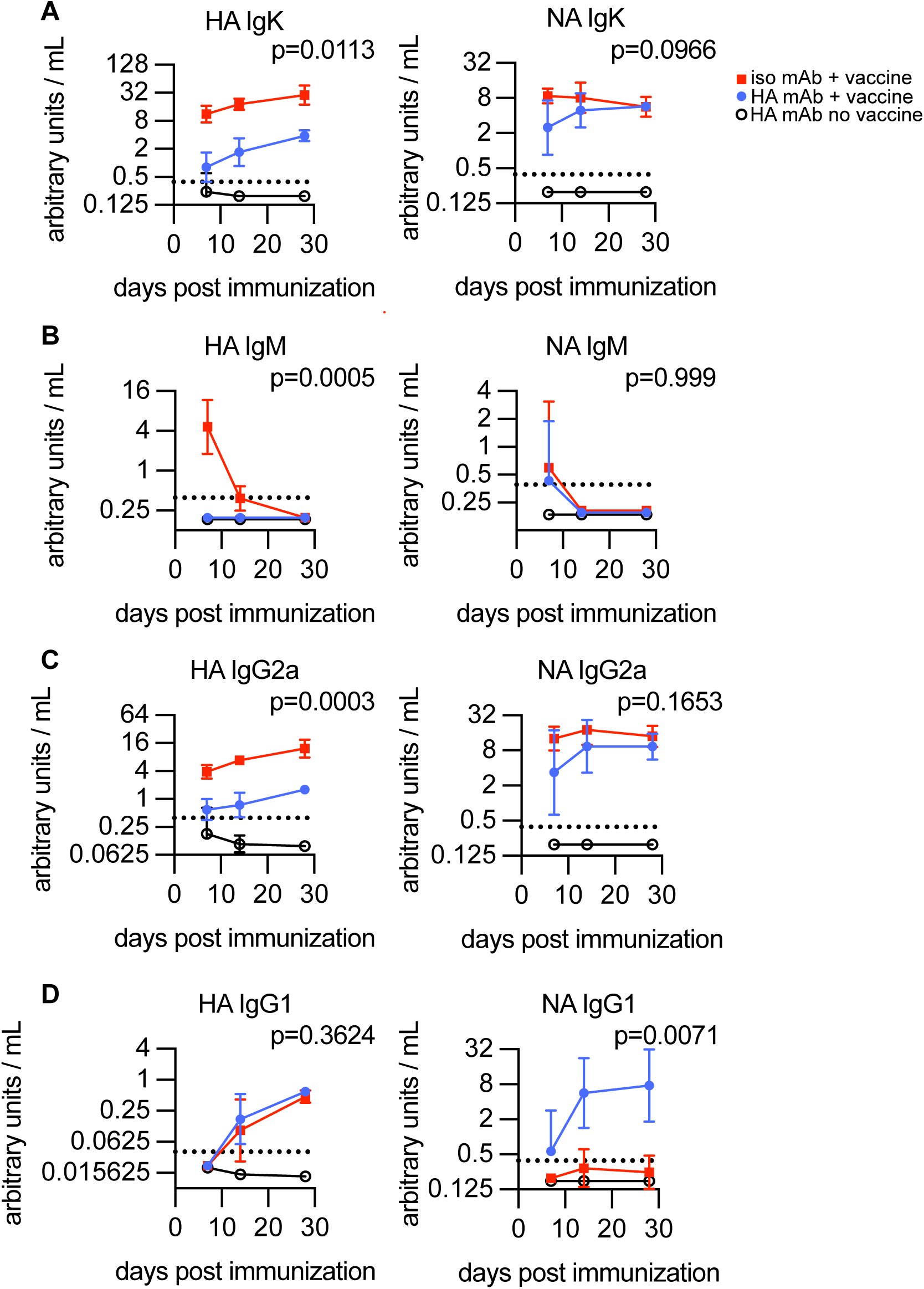
HA-specific antibody potently inhibits antibody responses to HA but not NA. BALB/c mice received either 1μg H28-D14 mAb (blue circles) or 1μg isotype control (red squares) prior to intramuscular immunization with 2,000 hemagglutinating units (HAU) of BPL-inactivated H1N1. Open circles represent serum from unimmunized mice that received 1μg H28-D14 mAb. IgK (A), IgM (B), IgG2a (C), and IgG1 (D) titers against HA and NA protein following immunization are shown. Dashed lines indicate the limit of detection. Data represents three independent experiments with n=3-5 mice per group. Geometric mean titer +/- 95% CI are plotted. Area under the curve (AUC) was calculated for individual mice and groups were compared with Welch’s ANOVA (A, C, HA D) with Dunnett’s T3 multiple comparisons or Kruskal-Wallis test (B, NA D) with Dunn’s multiple comparisons. Reported p-values for comparison between H28-D14 IgG2a (blue) and isotype control IgG2a (red) groups.

While mice with HA-specific mAb produced low levels of anti-HA antibody at 7 days post-vaccination, they failed to produce detectable levels of anti-HA IgM (**Fig. 1B**), suggesting that IgM responses are particularly susceptible to inhibition by circulating antibody. IgM responses to NA (**Fig. 1B**) at 7 days post-vaccination were low or undetectable in mice with and without HA mAb. The HA mAb also inhibited production of HA-specific, but not NA-specific, IgG2a (**Fig. 1C**), the dominant IgG subtype produced by BALB/c mice in response to influenza virus (Hovden et al., 2005). Conversely, and despite the overall reduction in anti-HA antibody, the mice with HA mAb produced IgG1 against HA and greatly enhanced IgG1 against NA (**Fig. 1D**). This implicates two distinct effects of circulating antibody on *de novo* antibody responses against influenza viruses: inhibition of antibody responses against the protein targeted by the antibody, and broad enhancement of IgG1 subclass switching.

### HA-specific mAb inhibits HA-specific B cells from becoming plasmablasts, germinal center B cells, and memory B cells

Next, we directly compared the ASC populations in the draining lymph nodes of mice with and without HA mAb at the time of vaccination. At 4 days post-vaccination, mice with HA mAb had significantly fewer plasmablasts than control mice, measured either by the fraction of B cells that were plasmablasts or total number of plasmablasts (**Fig. 2A, Supplementary Fig. 2A**). Mice with HA mAb also had fewer HA-specific plasmablasts (**Fig. 2B-C, Supplementary Fig. 2B**) as identified by binding to a fluorescently-labeled HA protein. At 7 days post-vaccination the overall plasmablast populations were similar between experimental groups (**Fig. 2A**), but the mice with HA mAb still had greatly reduced HA-specific plasmablasts (**Fig. 2B**). After 14 days post-vaccination, plasmablast numbers declined to background levels in draining lymph nodes.

**Figure 2:**
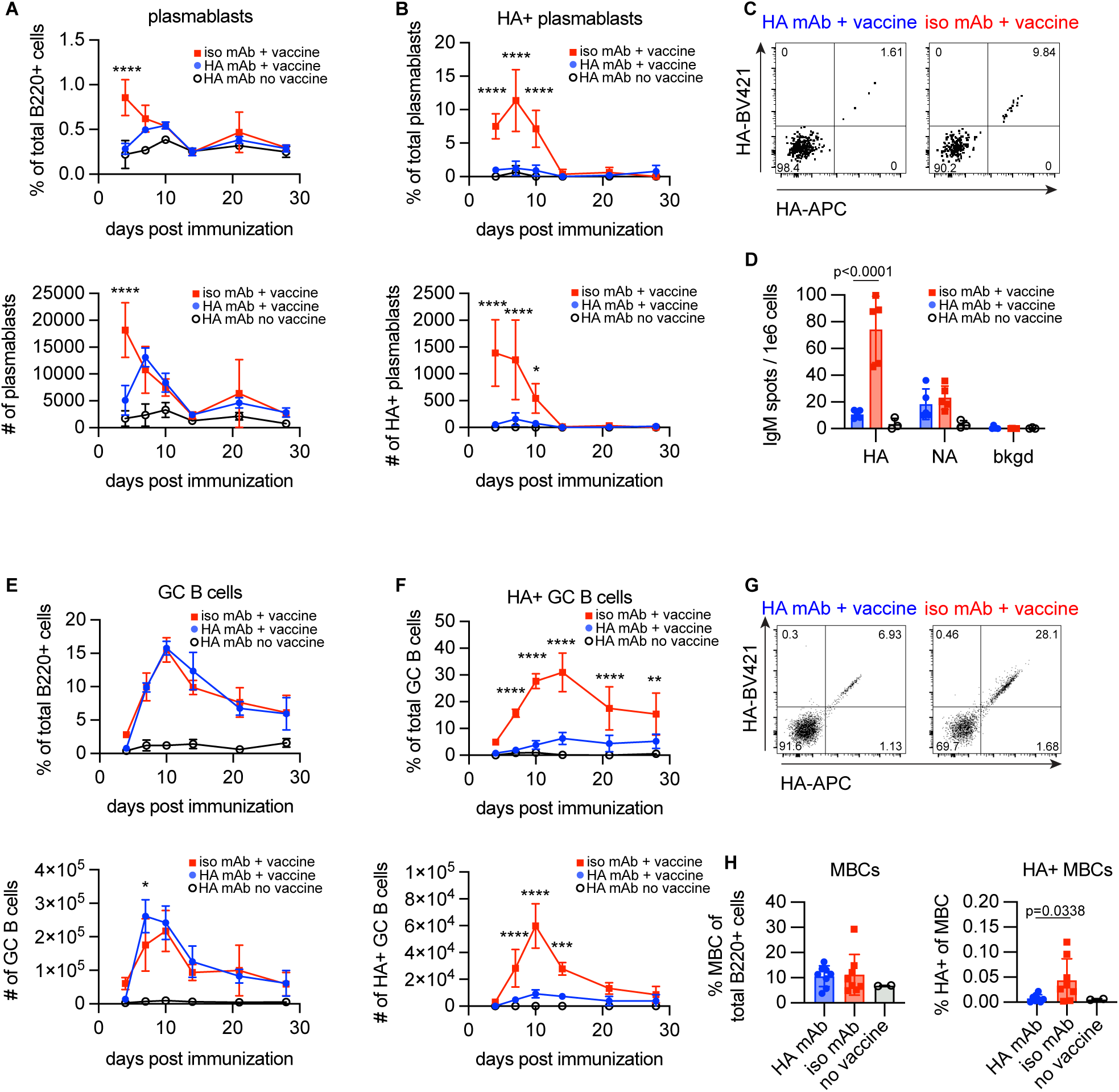
HA-specific antibody inhibits HA-specific B cells from becoming plasmablasts, germinal center B cells, and memory B cells. BALB/c mice received either 1μg H28-D14 mAb (blue circles) or 1μg isotype control (red squares) prior to intramuscular immunization with 2,000 HAU BPL-inactivated H1N1. Open circles represent unimmunized mice. (A) Plasmablasts (CD3^-^F4/80^-^Gr1^-^Ter119^-^B220^+^CD138^+^) and (B) HA probe-binding plasmablasts in draining lymph nodes of mice measured by flow cytometry at days 4, 7, 10, 14, 21, and 28 post-immunization. (C) Example flow plots of the plasmablast compartment at day 7 binding HA probe. (D) IgM-secreting cells in draining lymph nodes at day 7 post-immunization enumerated by ELISpot. (E) Germinal center B cells (CD3^-^ F4/80^-^Gr1^-^Ter119^-^B220^+^GL7^+^Fas^+^) and (F) HA probe-binding germinal center B cells in draining lymph nodes of mice measured by flow cytometry at days 4, 7, 10, 14, 21, and 28 post-immunization. (G) Example flow plots of the GC B cell compartment at day 14 binding HA probe. (H) Memory B cells (CD3^-^F4/80^-^Gr1^-^Ter119^-^B220^+^GL7^-^CD138^-^CD38^+^) and HA probe-binding memory B cells in the spleen measured by flow cytometry at >1 year post-immunization. Data represents four independent experiments with n=3-8 mice per group. Arithmetic mean +/- standard deviation are plotted. Dots (D, H) represent individual mice. Significant differences between H28-D14 IgG2a (blue circles) and ctrl IgG2a (red squares) groups by two-way ANOVA with Tukey’s multiple comparisons test (A, B, D-F) or unpaired T test (H) are reported. *p<0.05, ** p<0.01, *** p<0.001, **** p<0.0001, ns=not significant.

These data suggest that plasmablasts in mice with HA mAb were specific for viral proteins other than HA, such as NA. Since a fluorescent NA protein to quantify NA-specific plasmablasts by flow cytometry was not available, we enumerated HA- and NA-specific IgM-secreting cells 7 days after vaccination by ELISpot. Consistent with the serum and flow cytometry data, anti-HA IgM-secreting cells were reduced in mice with HA mAb, while anti-NA IgM-secreting cells were unaffected (**Fig. 2D**). Collectively, these data indicate the inhibitory effect of HA antibody on early plasmablast responses is limited to HA-specific cells.

We next evaluated the effects of HA mAb on germinal center formation. While some studies describe antibody-mediated epitope-specific inhibition of germinal center B cells (Forsell et al., 2017; Inoue et al., 2023), another study reported that maternally-transferred anti-HA antibody suppressed HA serum antibody responses without reducing the number of HA-binding germinal center B cells in mice (Vono et al., 2019). We measured germinal center responses throughout the first 4 weeks following vaccination and found that the HA mAb didn’t affect the overall magnitude of the germinal center response (**Fig. 2E**), but did considerably reduce the number of HA-specific germinal center B cells **(Fig. 2F-G)**, suggesting that the mAb inhibited germinal centers in an antigen-specific manner. Since by 7 days post-vaccination the total HA IgG2a levels in control mice exceed those in mice given HA mAb (**Fig. 1C**), these results suggest that anti-HA antibody present at the initial time of vaccination is a potent inhibitor of HA-specific germinal center responses.

We also measured the impact of HA-specific mAb on the long-lived memory B cell (MBC) population. MBCs are an essential component of immune memory and are thought to be the main drivers of OAS. They can be produced by germinal centers (Inoue & Kurosaki, 2023), although many differentiate outside germinal centers in the first few days following vaccination (Glaros et al., 2021). More than one year after vaccination, mice given HA mAb had fewer HA-specific memory B cells than controls, with little to no enrichment of HA-specific memory B cells over naïve counterparts (**Fig. 2H**). If these findings are broadly applicable, failure to generate a robust memory B cell population in the presence of circulating antibody may represent a key inferiority of secondary immune responses and be a major driver of OAS.

### FcγR interactions mediate increased IgG1 production but are dispensable for the inhibitory effects of HA antibody

Some mechanisms of antibody-mediated inhibition of immune responses rely on engagement of Fc-gamma receptors (FcγRs). For example, BCR binding of immune complexed antigens can cause ligation of the inhibitory Fc-gamma receptor (FcγRIIB), which antagonizes intracellular signaling by the BCR (Muta et al., 1994) and immune synapse formation (Liu et al., 2010). Some studies implicate engagement of this receptor in antibody-mediated inhibition of *de novo* B cell responses (Kim 2011). Another proposed mechanism of inhibition is enhanced antigen clearance by antibody opsonization of antigen via antibody-Fc receptor interactions.

To assess if FcγRs were involved in our experimental system, we generated a mutated version of the HA mAb (a D265A substitution in the heavy chain) that cannot bind to FcγRs (Baudino et al., 2008). This mutation does not affect binding to the neonatal Fc receptor (FcRn) so the half-life of this mAb is unaltered. The HA mAb with the D265A substitution inhibited IgK, IgM, and IgG2a responses against HA but had no impact on production of these antibody types to NA (**Fig. 3A-C).** This antibody also reduced HA-specific, but not total, plasmablast and germinal center B cells (**Supplementary Fig. 3A-C**). Unlike the wild type HA mAb, the HA mAb with the D265A substitution did not increase IgG1 titers to NA (**Fig. 3D**). These results indicate that the inhibitory effects of the HA mAb are not mediated through FcγR interactions, while the broad enhancement of IgG1 requires mAb-FcγR interactions. The HA IgG1 response may represent the intersection of these opposing effects, where the general inhibition of HA antibody and the promotion of IgG1 class switching combine to produce a net null or modest effect. The mechanism for IgG1 enhancement in our experiments remains undefined, but likely involves changes in the polarization of CD4 T cells, which mediate antibody class switching and subtype selection (Stavnezer, 1996). Consistent with this, antibody-antigen immune complexes have been observed to promote Th2 differentiation of CD4 T cells and IgG1 production in mice (Anderson & Mosser, 2002; Bandukwala et al., 2007).

**Figure 3:**
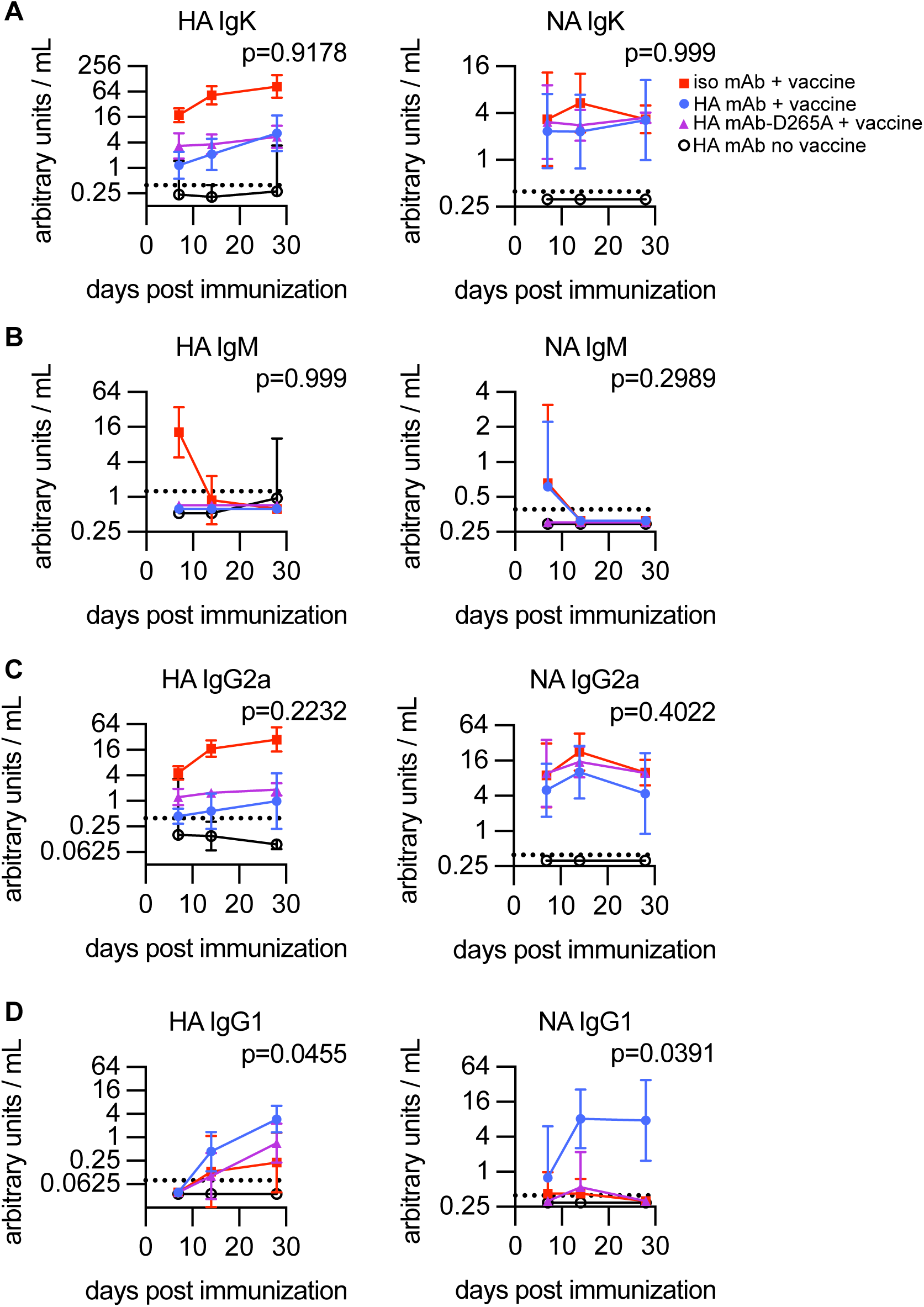
FcγR interactions mediate IgG1 increase but are dispensable for the inhibitory effects of HA antibody. BALB/c mice received either 1μg H28-D14 mAb (blue circles), 1μg H28-D14 with D265A mutation (purple triangles), or 1μg isotype control (red squares) prior to intramuscular immunization with 2,000 HAU BPL-inactivated H1N1. Open circles represent unimmunized mice that received 1μg H28-D14 mAb. Serum IgK (A), IgM (B), IgG2a (C), and IgG1 (D) titers against HA and NA protein following immunization are shown. Dashed lines indicate the limit of detection. Data represents three independent experiments with n=3-5 mice per group. Geometric mean titer +/- 95% CI are plotted. Area under the curve (AUC) was calculated for individual mice and groups were compared with Welch’s ANOVA (A, C, HA D) with Dunnett’s T3 multiple comparisons or Kruskal-Wallis test (B, NA D) with Dunn’s multiple comparisons. Reported p-values for comparison between H28-D14 IgG2a (blue circles) and H28-D14 D265A (purple triangles) groups.

### Anti-HA antibody inhibits de novo antibody responses to multiple HA epitopes, but most strongly epitopes shielded by the transferred antibody

Given that FcγR interactions are not required for HA mAb inhibition of *de novo* HA antibody responses, it is possible that the transferred mAb sterically prevents binding of antigen-specific B cells through epitope masking. If so, the inhibitory effect of the HA mAb should decrease as distance from the mAb’s cognate epitope increases. To assess this, we measured the *de novo* antibody response to the five defined antigenic sites in the head domain of the H1N1 protein: Sa, Sb, Ca1, Ca2, and Cb. The HA mAb used in our studies binds to the Sb site, and is nearest to the Sa, Ca1, and Ca2 sites, and farther from the Cb site (Caton et al., 1982; Gerhard et al., 1981) (**Fig. 4A**). We measured serum antibodies directed against each of the five HA antigenic sites using HAs that were engineered to retain only a single native antigenic site (**Supplementary Fig. 4**) (Angeletti et al., 2017). As a control, we also measured antibody binding to an HA (S12a) that possesses mutations in all five of the major antigenic sites (Das et al., 2013).

**Figure 4:**
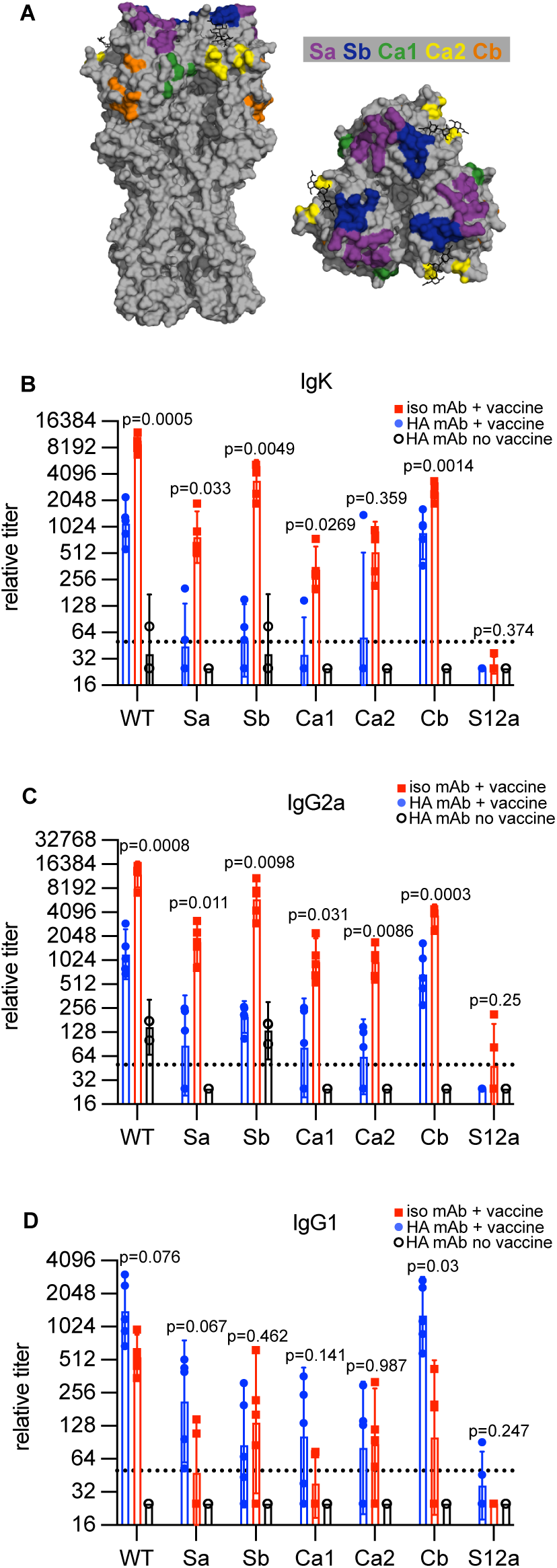
Anti-Sb antibody most potently inhibits *de novo* antibody responses to Sb antigenic site and nearby neighboring antigenic sites. (A) Structure of PR8 H1N1 HA protein with residues contributing to each antigenic site indicated by color. (B-D) BALB/c mice received either 1μg H28-D14 mAb (blue circles) or 1μg isotype control (red squares) prior to intramuscular immunization with 2,000 HAU BPL-inactivated H1N1. Open circles represent serum from unimmunized mice that received 1μg H28-D14 mAb. Serum IgK (B), IgG2a (C), and IgG1 (D) titers against PR8 HA epitope panel at day 28 post-immunization are shown. Data points represent individual mice. Dashed lines indicate the limit of detection. Data represents three independent experiments with n=3-5 mice per group. Geometric mean +/- 95% CI are plotted. Significant differences between H28-D14 (blue circles) and control IgG2a (red squares) groups by unpaired t-test with Welch’s correction are reported.

Control mice that received a non-influenza mAb produced antibodies to all five of the antigenic sites, while mice who received HA mAb produced lower amounts of antibodies to each of the HA antigenic sites (**Fig. 4B**). The Sb mAb strongly inhibited *de novo* responses to the Sb antigenic site and to the neighboring Sa, Ca1, and Ca2 antigenic sites. *De novo* responses to the Cb antigenic site, the farthest from the Sb antigenic site, were minimally affected by the passively transferred Sb mAb. The Sb mAb had varying effects on the production of individual IgG subtypes. The Sb mAb inhibited IgG2a production against each antigenic site, with greater inhibition of responses to antigenic sites near the Sb antigenic site (**Fig. 4C**). Conversely, the Sb mAb did not inhibit IgG1 production against the Sb, Sa, Ca1, and Ca2 antigenic sites, and promoted IgG1 production against the distant Cb antigenic site (**Fig. 4D**). Collectively, these data demonstrate that the inhibitory effect of the anti-HA antibody in our experiments is less pronounced against antigenic sites that are farther from the binding site of the mAb, with enhancement of IgG1 production against more distant antigenic sites.

## Discussion

The mechanism and extent of antibody-mediated inhibition of *de novo* immune responses has been well studied. Much of this work has evaluated models that involve Fc-mediated effects and epitope masking. Data in support or opposition of epitope masking can, as described by Heyman and colleagues (Heyman, 1999; Xu & Heyman, 2020), be broadly grouped into two categories: 1. the necessity of Fc-mediated phenomena (such as Fc-receptor signaling, antigen clearance, and complement binding), and 2. the antigenic breadth of inhibition.

Many studies have used F(ab’)2 fragments to interrogate the involvement of the Fc-region, with some reporting no change in inhibition (Cerottini et al., 1969; Karlsson et al., 1999; Tao & Uhr, 1966) and others observing markedly reduced inhibition (Dangi et al., 2023; Enriquez-Rincon & Klaus, 1984; Sinclair, 1969; Sinclair et al., 1968). However, F(ab’)2 fragments are an imperfect surrogate for Fc-null antibody, as their shorter half-life and smaller size may reduce their capacity to mask epitopes. In our studies, a full-size HA mAb with the D265A substitution that eliminates FcγR binding inhibited *de novo* anti-HA antibody responses. These results align with other studies that demonstrate that antibodies can inhibit B cell responses in FcγR-knockout mice (Bernardo et al., 2015; Karlsson et al., 1999). The D265A substitution does not fully eliminate all Fc-mediated interactions; these antibodies can still bind complement and engage non-canonical Fc receptors that bind the glycosylated Fc region including DC-SIGN, CD23, and FCRL5 (Jennewein & Alter, 2017; Wang et al., 2015). While it remains possible that the HA mAb in our experiments inhibits through such interactions, the limited antigenic breadth of the inhibitory effect suggests otherwise.

Unlike the inhibitory effects, the antibody-mediated enhancement of IgG1 responses was most pronounced for the non-HA vaccine antigen, NA, and involved signaling through FcγRs. Other groups have observed that IgG1 responses to a haptenated protein antigen were increased when delivered with an anti-hapten antibody (Coulie & van Snick, 1985; Wernersson et al., 2000; Wiersma et al., 1989). Some of these studies found that immune complexes with IgG2a antibodies (the subtype used in our studies) but not IgG1 antibodies increased IgG1 production. These results are consistent with an FcγR-dependent effect, as IgG1 and IgG2a have different interaction profiles with FcγRs: IgG2a binds better to activating FcγRs and IgG1 binds better to the inhibitory FcγRIIB (Nimmerjahn & Ravetch, 2014). Future studies should evaluate if passively transferred IgG1 HA mAbs affect immune responses differently compared to IgG2a HA mAbs.

Previous studies have shown that an anti-Sb Fab narrowly inhibits Sb responses with little inhibition of responses to neighboring antigenic sites (Angeletti et al., 2017). In our experiments, we passively transferred a full-length anti-Sb mAb, and observed potent inhibition of *de novo* responses against the Sb antigenic site, as well as inhibition of responses against other nearby antigenic sites. Collectively, this suggests that a fully intact HA antibody can occlude—and thereby exert its inhibitory function on—neighboring epitopes that it does not directly contact. This is consistent with observations that Sb antibodies and antibodies to neighboring sites (e.g. Sa, Ca) cannot concurrently bind to HA from detergent-treated virus (Lubeck & Gerhard, 1981). The steric constraints are likely greater *in vivo*, where soluble antibody interferes with membrane-bound BCRs binding to antigen on the surface of virions. It will be important for future studies to evaluate the inhibitory potential of mAbs that recognize other HA epitopes. Antibodies that bind to epitopes that are lower on HA, such as Cb or the stalk, may not have reciprocal inhibitory effects on epitopes higher on the HA globular head, including Sa and Sb. Differences in vaccine format, such as changes in antigen density and display (i.e. membrane bound versus soluble), may also affect epitope accessibility and thereby alter the inhibitory effect of antibodies.

Although the circulating antibody in our studies was monoclonal, it inhibited 80-90% of the *de novo* HA antibody response. While circulating antibody most strongly inhibited *de novo* antibodies of similar specificities, we found no net increase of total antibody responses to other epitopes. These findings suggest that, in the context of influenza vaccination, pre-existing antibody to a conserved epitope could have a substantial inhibitory effect on *de novo* responses to neighboring novel epitopes. Therefore, circulating antibody present at the time of vaccination might affect the overall magnitude and specificity of antibodies against novel epitopes.

Beyond the impact on *de novo* antibody production, we observed that passively transferred HA mAb reduced HA-specific B cells in the early plasmablast, germinal center, and, perhaps most significantly, the memory B cell (MBC) response. If circulating antibody prevents the formation of a robust MBC population, then MBCs induced in the absence of antibody (i.e., from a primary exposure) are expected to be more abundant than MBCs generated in subsequent exposures. This dynamic is consistent with the dominance of primary-response-elicited antibodies characteristic of OAS. Even a more subtle impact on MBC development— such as reduced MBC diversity or longevity rather than elimination—might account for year of birth related differences in influenza virus susceptibility (i.e. imprinting) (Arevalo et al., 2020; Gostic et al., 2016).

Since all of our passive transfer experiments were completed in naïve mice, we only examined how antibodies affect naïve B cells, and we did not evaluate if antibodies affect already established MBCs. Other groups have observed that antibodies can inhibit activation of MBCs in an antigen-specific manner (McNamara et al., 2020; Pape et al., 2011), but the relative inhibitory potency of antibody on naïve and MBCs remains to be determined. If the inhibitory effects of HA antibodies are less potent on MBCs, this would further lead to preferential recruitment of MBCs during secondary influenza virus encounters.

## Acknowledgements

This project has been funded in part with Federal funds from the National Institute of Allergy and Infectious Diseases, National Institutes of Health, Department of Health and Human Services, under Contract No. 75N93021C00015 (S.E.H.) and Grant No. 1R01AI108686 (S.E.H). S.E.H. holds an Investigators in the Pathogenesis of Infectious Disease Awards from the Burroughs Wellcome Fund. We thank Dr. Ed Behrens for guidance on statistical analyses.

## Author Contributions

Conceptualization, E.G. and S.E.H.; Investigation, E.G.; Formal Analysis, E.G.; Visualization, E.G. and S.E.H.; Writing – Original Draft, E.G., Writing – Review & Editing, E.G., L.C.E., and S.E.H.; Supervision, L.C.E. and S.E.H..; Funding Acquisition, S.E.H.; Resources, J.S.G. and J.W.Y.

## Methods

### Vaccine

PR8 H1N1 virus was generated through reverse genetics as previously described (Neumann et al., 2005) and then expanded in day 10 embryonated hens eggs (Charles River). At 48 hours post infection, allantoic fluid was harvested, clarified, and inactivated with 0.1% v/v beta-propiolactone (BPL) with 100mM HEPES for 16 hours at 4°C. The inactivated virus was then incubated at 37°C for 2 hours. Virus was concentrated via ultracentrifugation over a 20% w/v sucrose cushion for 1 hour at 20,000 rpm, then resuspended in phosphate buffered saline (PBS) and titered by a hemagglutination unit assay.

### Antibodies and proteins

Recombinant PR8 HA proteins, including the Y98F mutant HA used for flow cytometry and the delta4 variants fused for epitope mapping, were produced in-house in 293F cells. Recombinant PR8 NA protein was produced by Sino Biological. The H28-D14 mAb was produced by Biointron. Mouse IgG2a isotype control monoclonal antibody E23 was obtained from BEI Resources (NR-15540).

### Mice and immunizations

All experiments using mice were approved by the Institutional Animal Care and Use Committees at the Wistar Institute and the University of Pennsylvania. Female BALB/c mice were purchased from Charles River Laboratories. At 6-12 weeks of age, mice were injected intraperitoneally with 1μg of monoclonal antibody in 100μL of PBS. Six hours later, mice were immunized intramuscularly in both gastrocnemius muscles with a total of 2,000 HAU of BPL-inactivated PR8 H1N1 virus (approximately 1,000 HAU per leg). Blood was collected from mice by submandibular puncture at indicated time intervals into collection tubes (Sarstedt, 41.1378.005) from which serum was isolated and stored at −20°C.

### ELISAs

Immulon 4HBX 96-well plates (Fisher, 14-245-153) were coated with 50 μL protein antigens at 2μg/ml in PBS and incubated overnight at 4°C. Plates were washed three times and blocked for one hour with 125 μL of blocking buffer (PBS with 3% goat serum, 0.5% nonfat milk, 0.1% Tween 20). Two-fold serial dilutions of serum samples were made in blocking buffer starting at a 1:50 dilution. Plates were washed three times, then diluted serum was added to plates and incubated for two hours. Detection antibodies conjugated to HRP were diluted in blocking buffer: rabbit anti-mouse IgM (Jackson Immunoresearch, 315-035-049) diluted 1:5000; goat anti-mouse IgK (Novus, NB7549) diluted 1:5,000; goat anti-mouse IgG1 (Southern Biotech, 1071-05) diluted 1:5,000; goat anti-mouse IgG2a (Southern Biotech, 1081-05) diluted 1:2,500. Plates were washed three times and diluted detection antibodies were added to plates and incubated for one hour. Plates were washed three times and 50μL of room temperature KPL SureBlue TMB substrate was added to wells. Plates incubated for 5 minutes before quenching with 25μL of 250mM HCl. Well absorbance was read at 450 nm. All wash steps were conducted using PBS with 0.1% Tween 20.

### B cell ELISpot

Plates (MilliporeSigma, S2EM004M99) were activated with 50 μL of 35% ethanol and washed 4 times with PBS. 100 μL of 10μg/mL of protein antigens was applied to wells and plates were incubated overnight at 4°C. Before cells were added, plates were washed 4 times with PBS and blocked with RPMI 10% FBS at 37°C for 1 hour.

For IgM ELISpot, mice were sacrificed 7 days post-immunization and popliteal, inguinal, and iliac lymph nodes were collected. Lymph nodes were dissociated by grinding with frosted microscope slides and rinsed with PBS through a 70 um filter. Cells were washed twice with PBS, resuspended in RPMI 10%FBS, and added to plates. Plates incubated overnight at 37°C. To develop, cells were discarded and plates washed 4 times with PBS 0.1% Tween 20. Rabbit anti-mouse IgM HRP (Jackson Immunoresearch, 315-035-049) was diluted 1:5000 in PBS 10% FBS 0.1% Tween and added to wells. Plates incubated at 37°C for 2 hours, then were washed 3 times with PBS 0.1% Tween and 3 times with PBS. AEC substrate (BD Biosciences, 551951) was added to wells and plates developed at room temperature until spots were visible, approximately 30 minutes. Plates were rinsed 5 times with DI water and set out to dry. Well images were captured with Immunospot (S6 Universal M2) and spot numbers were enumerated by hand.

### Flow cytometry

For analysis at days 4-28 post-immunization, mice were sacrificed and popliteal, inguinal, and iliac lymph nodes were collected. Lymph nodes were dissociated by grinding with frosted microscope slides and rinsed with PBS through a 70 μm filter. Cells were washed twice with PBS and incubated for 20 minutes on ice with 1:100 dilution anti-CD16/32 (clone 2.4G2, BD biosciences) and 1:500 dilution Live/Dead nearIR dye (ThermoFisher). Cells were washed twice with PBS 1% FBS 2mM EDTA and incubated with fluorescent antibodies listed below and HA probes on ice for 1 hour. Cells were washed and resuspended in PBS 1% FBS 2mM EDTA and run on a Cytoflex LX (Beckman Coulter).

For memory B cell analysis, mice were sacrificed and spleens were collected. Spleens were dissociated by grinding with frosted microscope slides and rinsed with PBS through a 70 μm filter. Cells were resuspended in 2 mL ACK lysis buffer (Gibco) and incubated for 90 seconds for erythrocyte rupture before addition of 48 mL of PBS. Cells were pelleted, resuspended, and strained again through a 70 μm filter prior to staining as described above.

To reduce non-specific binding of the HA probe via interactions with cell-surface sialic acid, recombinant PR8 HA protein was produced with a Y98F mutation. Purified HA was biotinylated using the BirA500 kit (Avidity) and buffer exchanged into PBS with a 30kDa spin column (MilliporeSigma) to remove excess biotin. Prior to incubation with cells, HA protein was incubated with streptavidin-APC (Biolegend, 405207) and streptavidin-BV421 (Biolegend, 405225) independently at a molar ratio of 1:4 streptavidin:HA monomer (for an approximately 1:1 ratio of binding sites) for 20 minutes on ice. HA proteins were then added to the antibody staining mix at a final concentration of 50nM HA trimer for each fluorophore (final total HA trimer concentration 100nM).

Antibody staining mix contained antibodies against CD38-AF700 (Biolegend, 102742), CD138-BB515 (BD, 564511), B220-BV605 (Biolegend, 103244), Ter119-PE/Cy7 (Biolegend, 116222), F4/80-PE/Cy7 (Biolegend, 123114), CD3-PE/Cy7 (Biolegend, 100220), Gr1-PE/Cy7 (Biolegend, 108416), GL7-PerCP/Cy5.5 (Biolegend, 144610), and Fas-PE (BD, 561985).

