## Supplemental figures for "Influenza virus antibodies inhibit antigen-specific *de novo* B cell responses in mice"

### Goodwin et al. 2024 Supplemental Figure 1

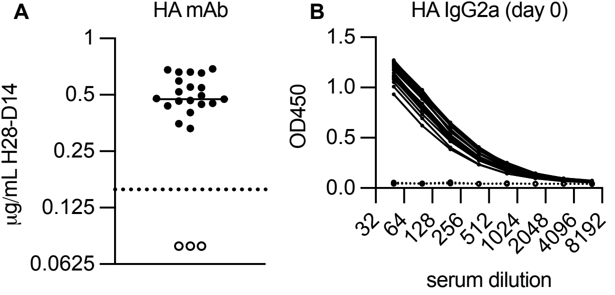

**Supplementary figure 1**

(A) Antibody titers of mice given 1ug H28-D14 mAb (black dots) or no mAb (open circles) intraperitoneally 5 hours prior to serum collection calculated from ELISA values of serially diluted sera (shown in B). Data points represent individual mice.

### Goodwin et al. 2024 Supplemental Figure 2

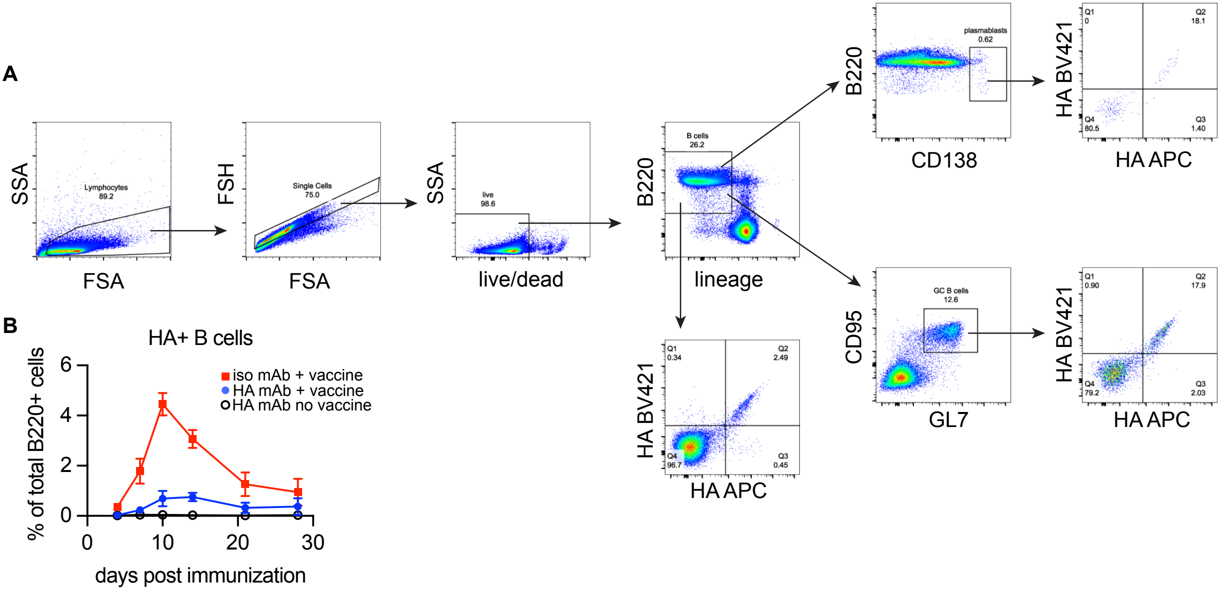

**Supplementary figure 2**

(A) Representative flow gating plots for plasmablasts and germinal center B cells, from an isotype control mouse at day 7 post-immunization. (B) Total B cells ($CD3^{-}F4/80^{-}Gr1^{-}Ter119^{-}$ $B220^{+}$) that bind the HA probe in draining lymph nodes at days 4, 7, 10, 14, 21, and 28 post-immunization, from the same mice shown in figure 2. Significant differences between H28-D14 IgG2a (blue) and ctrl IgG2a (red) groups by two-way ANOVA with Tukey's multiple comparisons test are shown. * $p<0.05$, ** $p<0.01$, *** $p<0.001$, **** $p<0.0001$.

### Goodwin et al. 2024 Supplemental Figure 3

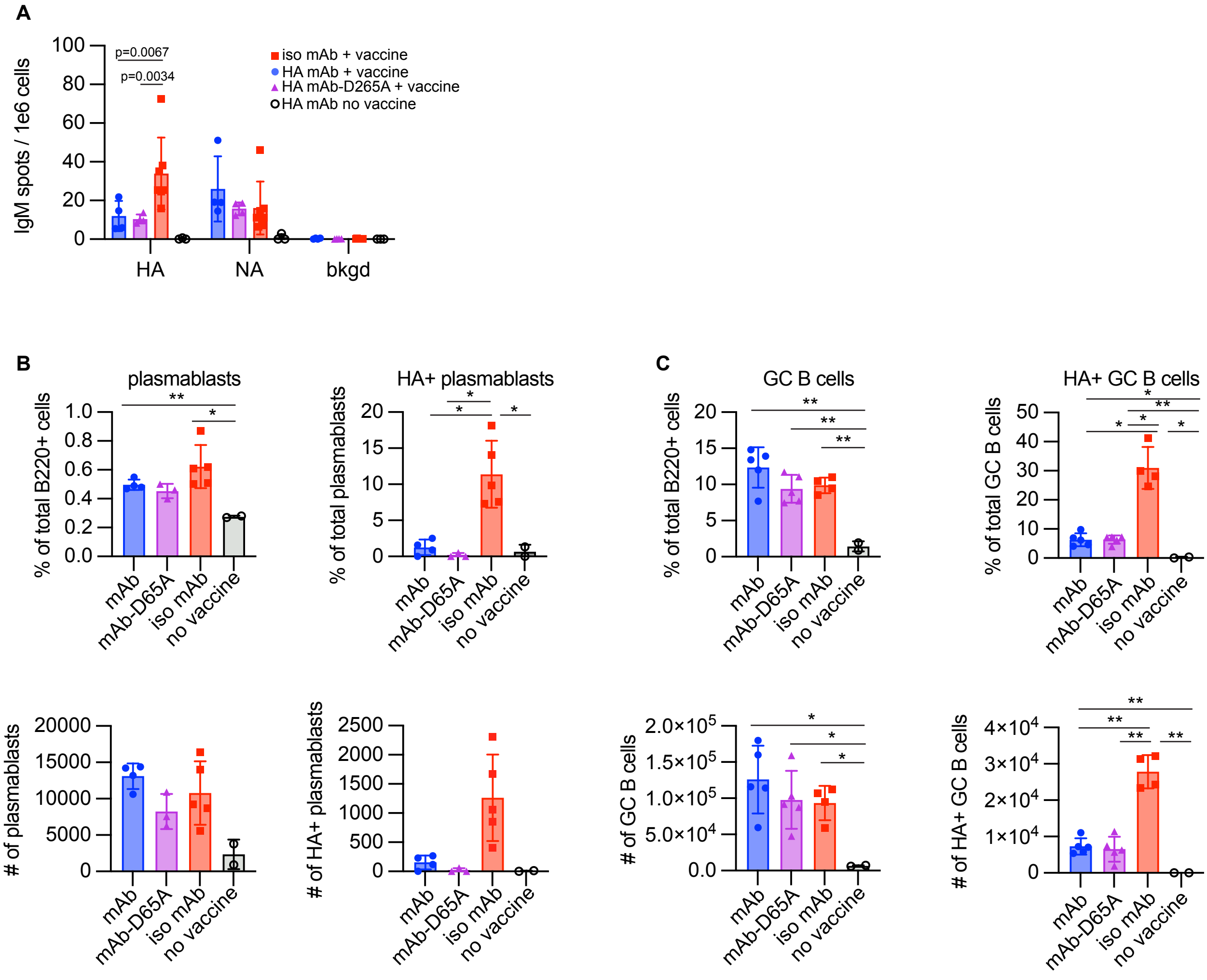

**Supplementary figure 3**

(A) IgM-secreting cells in draining lymph nodes at day 7 post-immunization enumerated by ELISpot. (B) Plasmablasts ($CD3^{-}F4/80^{-}Gr1^{-}Ter119^{-}B220^{+}CD138^{+}$) and HA probe-binding plasmablasts in draining lymph nodes of mice measured by flow cytometry at day 7 post-immunization. (C) Germinal center B cells ($CD3^{-}F4/80^{-}Gr1^{-}Ter119^{-}B220^{+}GL7^{+}Fas^{+}$) and HA probe-binding germinal center B cells in draining lymph nodes of mice measured by flow cytometry at day 14 post-immunization. Data represents two independent experiments with n=3-5 mice per group. Data points represent individual mice. Arithmetic mean +/- standard deviation are plotted. Significant differences determined by Welch ANOVA with Dunnett's T3 multiple comparisons test are indicated. * $p<0.05$, ** $p<0.01$, *** $p<0.001$, **** $p<0.0001$.
